# Isolation and metabolomic analysis of the culturable human gut mycobiota during dysbiosis

**DOI:** 10.64898/2026.01.28.702060

**Authors:** Yanet Teresa Cruz, Adán Alcántara Reyes, Patricia Espinosa Cueto, Vanessa Ruíz-Villegas, Albert D. Patiño, Mario Figueroa, Alba Romero Rodríguez

## Abstract

The human gut mycobiome, though less diverse than the bacterial microbiome, plays a significant role in health and disease. This study investigates the culturable fungal communities in fecal samples from hospitalized patients with diarrhea in Mexico City. We isolated and characterized 26 fungal strains using culture-dependent methods, including 20 yeasts and six filamentous fungi. The most prevalent organisms were *Candida albicans, Rhodotorula mucilaginosa, Penicillium* spp., *and Paecilomyces* spp.

Fungal isolates were tested for their ability to withstand gut-like conditions, including temperature, pH, oxidative stress, and bile salts. Notably, *Paecilomyces variotii* demonstrated thermotolerance, surviving at 42°C, and exhibited competitive growth against other fungi. Co-occurrence analysis revealed associations between fungal isolates and bacterial pathogens such as *Salmonella* and *Clostridioides difficile*, suggesting potential interkingdom interactions.

Cytotoxicity assays on Caco-2 cells showed that cell-free supernatants from *Candida inospicua* and filamentous fungi reduced cell viability by up to 40%. Finally, dereplication and untargeted metabolomic analyses of P. variotii, Penicillium crustosum, and Penicillium chrysogenum revealed the presence of several bioactive metabolites, including mycotoxins and antimicrobial compounds, highlighting their potential roles in gut ecology and disease.

Overall, this study underscores the importance of the gut mycobiome in dysbiosis and its interactions with bacterial pathogens. The findings suggest that fungi, particularly thermotolerant species such as P. variotii, may contribute to gut dysbiosis and disease progression, particularly in immunocompromised patients. Further research is needed to elucidate the functional roles of these fungi and their metabolites in gut health and disease.

## Introduction

Fungal communities play critical ecological roles in terrestrial and aquatic environments, acting as decomposers, pathogens, and mycorrhiza (Zhang et al., 2020). Recent studies have shown that fungi associated with animals, particularly herbivores, are an essential component of the microbiome (Meili et al., 2023). Moreover, the human gastrointestinal tract is a thriving habitat for numerous fungal species, comprising 0.1% of the total gut microbes (Nash et al., 2017).

In contrast to bacterial communities, the human gut mycobiome exhibits limited diversity and high inter- and intra-variability. Over time, an individual’s mycobiome is no more similar to itself than it is to another person’s (Nash et al., 2017). Despite some limitations, including distinguishing true symbionts from transient fungi (Suhr and Hallen-Adams, 2015; Lavrinienko et al., 2021), several fungal species persist across many samples, indicating that a core gut mycobiome may exist. Ten genera are consistently found in the human gastrointestinal tract, including *Candida* species (mainly *Candida albicans*), *Saccharomyces*, *Penicillium*, *Aspergillus*, *Cryptococcus*, *Malassezia*, *Cladosporium*, *Galactomyces*, *Debaryomyces,* and *Trichosporon* (Wu et al., 2021; Ferrocino et al., 2022).

Understanding the functions of the mycobiome and its significance for intestinal homeostasis and disease pathogenesis is still in its infancy. It is proposed that gut fungi help train the immune system and contribute to overall gastrointestinal homeostasis (Kumamoto, 2016; Santus et al., 2021). However, colonization and growth of opportunistic fungal pathogens, as well as the expansion of resident fungi, can cause tissue damage through toxins and dysregulation of host immune responses, locally and distally, thereby influencing the disease course and outcome. Furthermore, several observations suggest that bacteria and fungi interact within the gut, influencing each other through different levels of symbiosis (Richard and Sokol, 2019). Consequently, modifications in a single factor, such as the bacteriome or the mycobiome, may impact the other. For instance, during antibiotic treatment, the bacterial component is dramatically affected, creating a permissive environment for the expansion of bacterial pathogens and pathobionts, such as *C. difficile* infection (CDI) (Mullish and Williams, 2018). Evidence suggests that CDI can be maintained or exacerbated by *C. albicans* (van Leeuwen et al., 2016; Lambooij et al., 2017), underscoring the role of fungi in the gut during bacterial infections.

Changes in fungal communities, accompanied by alterations in bacterial communities, can exacerbate dysbiosis [11], creating a tolerant environment for fungal expansion or colonization. With increasing numbers of patients needing intensive care, including older adults or immunosuppressed patients, infections caused by opportunistic and pathogenic fungi are growing. Strati *et al*. have suggested that commensal fungal species may initiate and promote the spread of specific diseases (Strati et al., 2016), implying that alterations in fungal communities are risky in healthy individuals. Likewise, alterations of the gut mycobiota have been linked to different pathologies, including metabolic disorders (obesity), Inflammatory Bowel Diseases (IBDs), and colorectal adenomas. Consequently, describing and isolating fungi in healthy individuals and during dysbiosis is crucial to understanding mycobiome composition and function. Also, fungi’s metabolic capacity should be evaluated to decipher its effect on the host and the intra- and interconnections between bacterial and fungal communities.

Although the “sequencing era” has significantly advanced our understanding of microbiome composition, culture-dependent methods remain essential for uncovering causal relationships and underlying mechanisms. In this study, we provide a comprehensive account of fungal cultivation techniques and phenotypic assays used to characterize the culturable gut mycobiota of patients with diarrhea in Mexico City. Additionally, we employed metabolomic analyses to investigate the metabolic capabilities of selected filamentous fungi.

## Methods

### Sample collection

This study was conducted using human fecal samples obtained as discarded biological material generated during routine hospital monitoring procedures at a hospital in Mexico City. The samples were transferred to the laboratory only after being designated for disposal. All samples were fully coded, and no direct or indirect patient identifiers or personal data were accessible to the researchers at any stage of the study. The research did not involve direct interaction with human subjects, nor the collection or use of identifiable private information.

### Isolation of cultivable fungal species from fecal samples

Stool samples were diluted in PBS and plated on Brain Heart Infusion agar (BHI BD Bioxon™) supplemented with antibiotics (rifamycin, 32 µg/mL; gentamicin, 16 µg/mL; apramycin, 32 µg/mL; chloramphenicol, 32 µg/mL; nalidixic acid, 32 µg/mL; cycloserin, 10 µg/mL; cefoxitin, 32 µg/mL; and ampicillin, 32 µg/mL) and incubated aerobically at 27°C for at least ten days with daily monitoring. Morphologically different yeast colonies and all molds were picked for further analysis. All selected isolates were further subcultivated to obtain pure colonies. Molds were subcultivated in Malt Extract Agar (agar, 15 g/L, malt extract, 40 g/L, peptone, 10 g/L) or Potato Dextrose Agar (agar, 15 g/L, dextrose, 20 g/L, potato extract, 4 g/L). Yeasts were subcultivated in YPD medium (10 g/L yeast extract, 20 g/L peptone, 20 g/L dextrose).

The isolates, grown on potato dextrose agar (PDA) or malt extract agar (MEA) for 48-72 h, were examined; colonial morphology and pigment production were recorded. The colony morphologies and pigmentation of the presumptive yeast isolates were also assessed on Candida chromogenic media (TM media™). The colors of each colony were recorded after 72 h of incubation.

### DNA extraction and Molecular characterization of fungal isolates

Genomic DNA from five- to seven-day-old cultures of filamentous fungi was extracted from either liquid SMA broth or PDA culture plates. The fungal biomass from the culture plates was scraped off, and the broth was obtained by vacuum filtration. Fungal biomass was desiccated by filtering and applying 1 ml of acetone. The dried mycelia were macerated in a mortar, and DNA extraction was conducted as previously described (Al-Samarrai and Schmid, 2000). Briefly, powdered mycelium was treated with lysis buffer (200 mmol/l Tris-HCl, 250 mmol/l NaCl, 25 mmol/l EDTA, 0·5% w/v SDS pH 8.5) and extracted with phenol: chloroform:iso-amyl alcohol (25:24:1). For strains refractory to phenol/chloroform extraction, DNA was obtained with the Quick-DNA™ Fungal/Bacterial Miniprep Kit (Zymo Research) according to manufacturer instructions.

For yeast, an isolated colony was cultured for 20-24 h at 30°C in YPD (10 g/L yeast extract, 20 g/L peptone, 20 g/L dextrose), and DNA was extracted as previously described (Harju et al., 2004).

Filamentous fungi strains were identified by amplifying and sequencing the ribosomal Internal Transcribed Spacer (ITS) region. The following primer set was used: ITS1 (5′-TCCGTAGGTGAACTTGC-3′) and ITS4 (5′-TCCTCCGCTTATTGATATGC-3′), as previously described (McCullough et al., 1998). The β-tubulin region was amplified using the primers Bt2a (5′-GGTAACCAAATCGGTGCTGCTTTC-3′) and Bt2b (5′-ACCCTCAGTGTAGTGACCCTTGGC-3′). For yeast, rDNA was amplified for each isolate targeting the ITS1-5.8S-ITS2 region, using the primers ITS5 (5′-GGAAGTAAAAGTCGTAACAAGG-3 ′) and ITS4 (5′-TCCTCCGCTTATTGATATGC-3′).

Sequences were assessed using Chromas Software (Version 2.6.6, Technelysium Pty. Ltd.), and their quality was assessed based on well-defined peaks for nucleotides in the chromatograms. Sequences were then analyzed by pairwise alignment against databases on the Micobank website (Robert et al., 2013). Fungal isolates were identified with a minimum of 97% sequence similarity and 95% coverage with a described species.

The MEGA software generated a maximum-likelihood phylogenetic tree from 1000 replicates. The phylogenetic tree was visualized and edited in MEGA.

### Phenotypical Characterization of Fungal Isolates

Three independent experiments were performed with yeast and filamentous fungi to assess the effect of oxidative stress, pH, and bile salts on fungal growth. Filamentous fungi were cultivated on MEA agar for 5-7 days. For each isolate, a 0.5 cm diameter agar plug was prepared. Mycelia plugs were placed in the center of each well of 24-well dishes containing SMA to evaluate the effects of oxidative stress (H2O2 at 2 and 5 mM), pH (4, 6, 6.8, and 8), and bile acids (taurocholate at 2 and 5 %). H_2_O_2_ was added to agar medium at ∼50◦C before solidification, and the medium was either used immediately or stored at 4◦C for no more than 24 h (Garrido-Bazan et al., 2018). Initial concentrations of H_2_O_2_ were 2 and 5 mM, but, as has been reported, H_2_O_2_ can react with medium components, and the exact concentration in the plates cannot be estimated (Garrido-Bazan et al., 2018). Growth was recorded from 72 to 120h.

### Interaction assays

Yeasts isolated from the same sample were cultivated in YPD media supplemented with 100 μg/mL gentamicin. A single yeast colony was transferred to YPD broth and incubated overnight. After incubation, 3μL of each yeast was mixed. After mixing the corresponding strains, 3 µL were plated in a YPD plate, and colony morphology was observed after 72 h.

Additionally, filamentous fungi and yeast from the same sample were cocultured. For this purpose, filamentous fungi were cultivated in YPD plates for five days or until sporulation was observed. Spores were collected in 500-700 μL of PBS buffer (for sporulation), and 3 µL of the spore suspension was mixed with 3 µL of yeast overnight culture. Plates were incubated at room temperature for seven days.

### Confrontation assays of filamentous fungi

For confrontation assays against *P. variotii* YTC22, filamentous fungi were cultivated on SMA for 7 days. At the end of the incubation time, a 0.5 cm plug of filamentous fungi was placed at one extreme, and on the other extreme, a 0.5 cm plug from a *P. variotti* YTC22 was inoculated. Plates were incubated at 29°C for 7 days. The test was done in triplicate [n=3]. Fungal growth inhibition was evaluated by measuring mycelial area using ImageJ.

### Cell culture and cell viability assay

Caco-2 cells (ATCC HTB-37 ™) were cultured in Dulbecco’s Modified Eagle Medium (DMEM12800-058, Gibco) containing 10% fetal bovine serum (FBS), 5 µg/ml penicillin-streptomycin, and 5 mM sodium pyruvate, and incubated at 37°C in a humidified atmosphere with 5% CO_2_. Caco-2 cells (1 × 10^4^) were seeded in 96-well plates and cultured for 3 days post-confluence for the cell viability assay. Cells were treated with cell-free supernatants or fungal organic extracts for 24 h. Cell viability was determined by 3-(4,5-dimethylthiazol-2-yl)–2,5-diphenyltetrazolium bromide (MTT) assay (Sigma-Aldrich, St. Louis, MO, USA). Cells were incubated with MTT reagent (5 mg/ml) at 37°C for 4 h. Subsequently, the purple crystals were resuspended with 100 μL of SDS-HCl solution at 37°C for 16h. Finally, the microplate was measured at 570 nm in a Multiskan FC Microplate Photometer (Thermo Fisher Scientific). The experiments were repeated in triplicate across at least 2 independent experiments, and the results were expressed as the average percentage of viable cells and compared with the control (untreated).

### Filamentous fungi culture and extracts

The strains *Paecilomyces variotii* YTC22, *Penicillium chrysogenum* YTC25, and *Penicillium crustosum* YTC26 were cultivated on 4 × PDA plates under aerobic conditions for 15 days at 37°C. Then, the mycelium for each condition was harvested from the plates and extracted with a 1:1 mixture of CHCl_3_: MeOH, shaking at 120 rpm for 2 h. The mixtures were filtered, and the filtrates were dried under vacuum. The organic extracts were stored at −4°C until use.

### Liquid chromatography tandem high**lZI**resolution mass spectrometry (LC**lZI**HRMS/MS), dereplication, and untargeted metabolomics analyses

The fungal organic extracts prepared at 1.0 mg/mL in 1:1 Dioxane-MeOH were subjected to LC-HRMS-MS/MS analysis (ESI positive mode) on a Q Exactive mass spectrometer (Thermo Fisher Scientific, Waltham, MA, USA) coupled with an Acquity ultraperformance liquid chromatography (UPLC) system (Waters Corp., Milford, MA, USA). The chromatographic conditions and MS parameters were set as described previously (Paguigan et al., 2017; El-Elimat et al., 2021). Dereplication of known metabolites was assessed by comparison of the fungal extracts data with known compounds’ retention times, *m/z* values, MS^2^ fragmentation patterns, and/or UV absorbance patterns from an *in house* dereplication library containing data for more than 750 authenticated standards (commercial and isolated mycotoxins) In addition, untargeted metabolomic analysis was performed via the Global Natural Products Social Molecular Networking (GNPS) platform (Wang et al., 2016; Aron et al., 2020; Nothias et al., 2020). For this, feature detection and processing of the HRMS-MS/MS data were performed using ProteoWizard (https://proteowizard.sourceforge.io/) and MZmine v3.8 (https://mzmine.github.io/download.html) using the following parameters: precursor ion mass tolerance: 0.02 Da; MS/MS fragment ion mass tolerance: 0.02; minimum matched peak: 6, and cosine score 0.7. Molecular networks were visualized with Cytoscape v.3.9.1 (Shannon et al., 2003). The annotation of compounds was at confidence levels 1 and 2, according to the Metabolomics Standards Initiative (mass accuracy <5 ppm) (Sumner et al., 2007).

## Statistical analysis

All raw data obtained throughout this study were processed and statistically analyzed. One-way ANOVA and two-sample t-tests were performed using GraphPad Prism V7.0 (San Diego, California, USA).

## Results

### Cultivable fungal isolates from fecal samples of patients with diarrhea

The cultivable mycobiota in 39 fecal samples from patients with diarrhea were investigated using selective media supplemented with antibiotics. Fungal growth was observed in approximately 40% of the samples, leading to the purification and subsequent identification of twenty yeast isolates (74% of the total) and six filamentous fungi (26%) (Fig. 1A). The taxonomic identity of the fungal strains was determined by molecular sequencing of the ITS rDNA region, followed by BLAST searches and maximum likelihood phylogenetic analysis. For certain filamentous fungi, β-tubulin gene sequencing was additionally performed to improve resolution. Yeast typing was conducted using ITS sequencing and chromogenic agar (Fig. 1B and Supporting Information).

**Figure 1.**
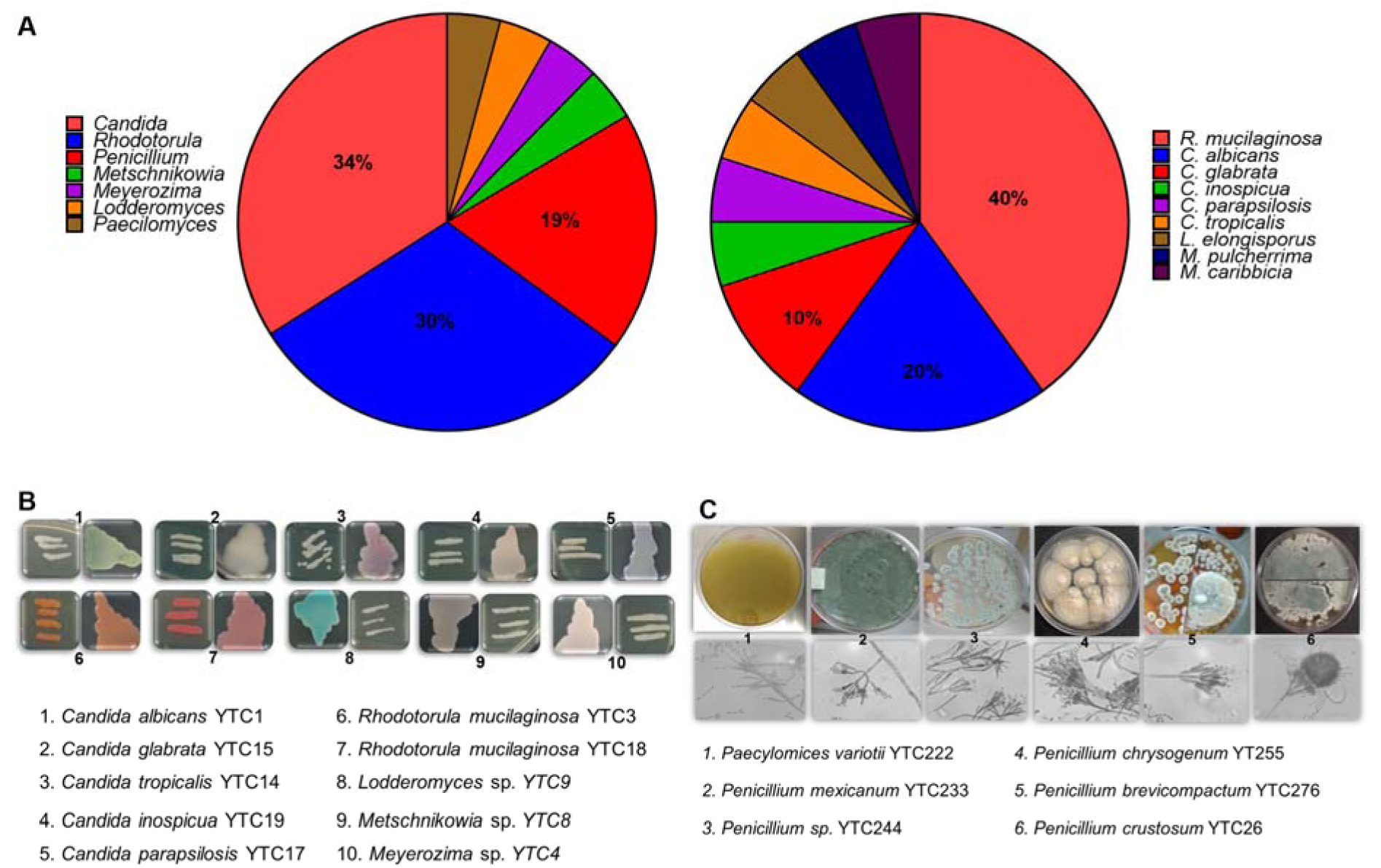
Culturable mycobiota in diarrheic samples of hospitalized patients. A) Pie charts of the percentage of fungal genera and species identified in fecal samples. B. Growth of fungal colonies in agar medium and yeast chromo agar. D. Colony morphology and microscopic characteristics of filamentous fungi.

Among the yeast isolates, *Candida* and *Rhodotorula* were the most prevalent genera (Fig. 1A; Fig. S1). Specifically, *C. albicans* and *Rhodotorula mucilaginosa* each accounted for 25 % of all yeast isolates (Fig. 1A). Less common *Candida* species, including *Nakaseomyces glabratus* (formerly *Candida glabrata*), *Candida inopsicua*, *Candida parapsilosis*, and *Candida tropicalis*, were each recovered from 5 % of the samples (Fig. 1B). In addition, we detected *Lodderomyces elongisporus*, an infrequent pathogen whose true incidence may be underreported due to its close genetic similarity to *C. parapsilosis* (Wang & Xu, 2023). Notably, potential probiotic yeasts from the *Meyerozyma* and *Metschnikowia* genera were also isolated (Fig. 1B).

For filamentous fungi, amplification and sequencing of the ITS and β-tubulin loci enabled the identification of the Penicillium and Paecilomyces genera (Fig. 1C and Supporting Information). Within *Penicillium*, five distinct taxa were detected: *Penicillium mexicanum* YTC23, *P. chrysogenum* YTC25, *P. crustosum* YTC26, and an unclassified *Penicillium* sp. YTC24, and *P. brevicompactum* YTC27. In the *Paecilomyces*, only *P. variotii* YTC22 was recovered.

### Intra- and inter-kingdom relationships

Polymicrobial interactions are increasingly recognized as critical determinants of disease outcomes, yet our understanding of how co-occurring microorganisms interact remains limited. To address this, fecal samples were screened using the FilmArray™ Gastrointestinal Panel, a multiplex PCR assay that detects 22 enteric pathogens, including *Salmonella* spp., *Escherichia coli*, and *Clostridioides difficile*. These molecular results were then correlated with culture-dependent fungal analyses. Notably, samples positive for *Salmonella* spp. and *E. coli* also yielded both yeast and filamentous fungi upon cultivation (Fig. 2A). Consistent with previous reports, *C. difficile* and *Candida albicans* were co-isolated from the same specimen (van Leeuwen et al., 2016; Brunetti et al., 2021). Additionally, *P. variotii* YTC22 was recovered from a *C. difficile*–positive sample (Fig. 2A).

**Figure 2.**
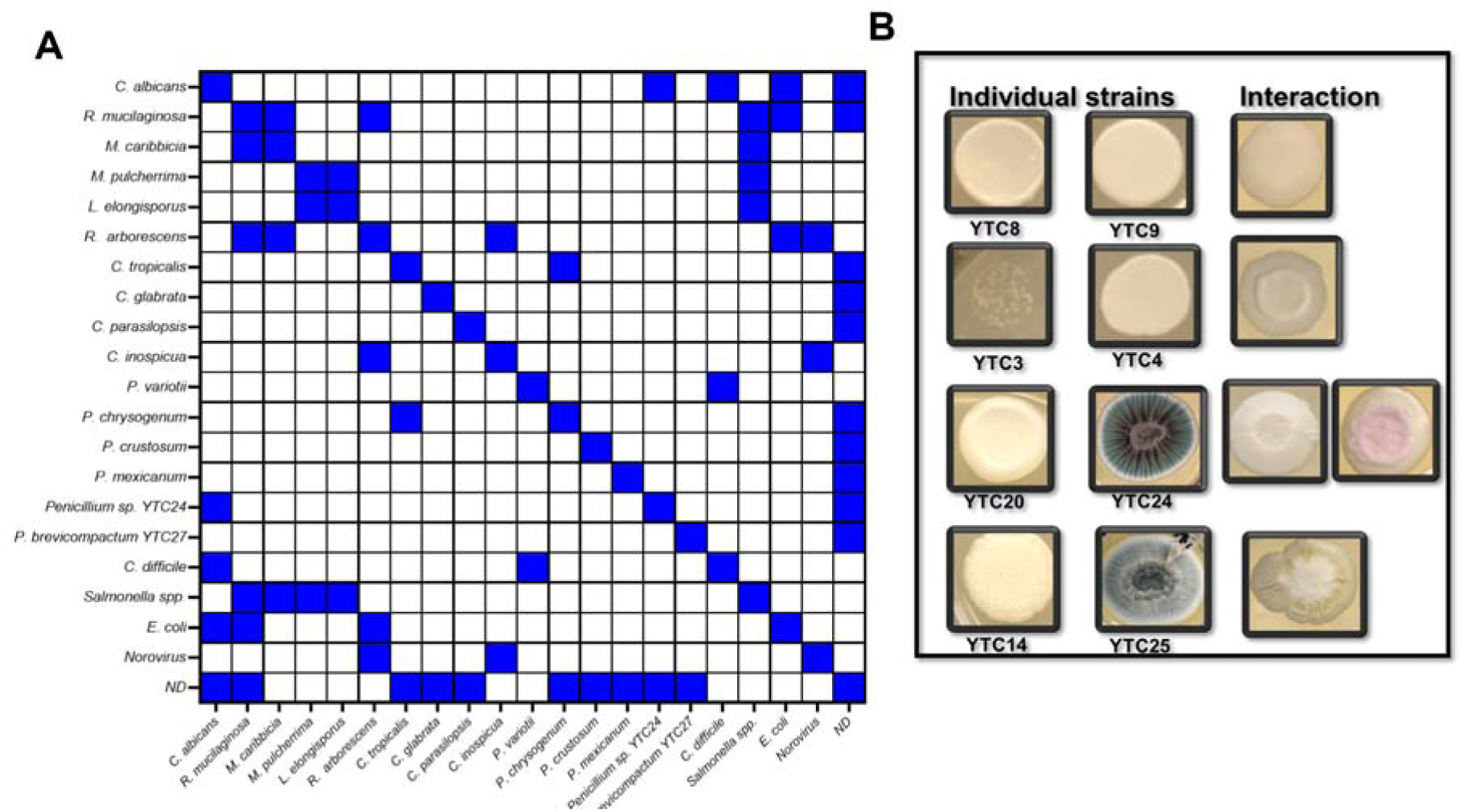
Intra and interkingdom occurrences. A. Matrix of co-occurrence based on panel array results and fungi isolation. B. Pairwise interactions between fungi isolated from the same donor. Pictures were taken 24 h after incubation at 30 °C.

Among samples positive for *Salmonella* spp., four yeast species, including *L. elongisporus*, were recovered (Fig. 2A). In norovirus-positive specimens, only *R. mucilaginosa* and *C. inopsicua* were cultivated. Interestingly, the majority of *Candida* spp. and all filamentous fungi were isolated from samples in which the FilmArray™ panel detected no bacterial, viral, or parasitic pathogens. No fungal growth was observed from samples positive for parasitic infections (Fig. 2A).

As previously noted, approximately 70 % of samples yielded a single fungal colony, either yeast or filamentous. In contrast, the remaining samples harbored multiple coexisting species (Fig. 2). To explore potential interactions, we established pairwise cocultures of fungal strains isolated from the same specimen. We monitored colony architecture for emergent features, which can signal coexistence dynamics (Gastélum et al., 2024). We also assessed chemical alterations in filamentous isolates grown alongside a partner strain on solid media. In the YTC8–YTC9 pair, colony morphology remained unchanged compared to each strain alone (Fig. 2B). In contrast, the YTC3–YTC4 interaction produced a colony morphology distinct from either yeast monoculture. For the YTC10–YTC11 duo, no novel architecture emerged, although both characteristic pigments were present (data not shown).

The interaction assay between the yeast and filamentous fungi resulted in the filamentous fungi growing on the top of the yeast colony (Fig.2b). In comparison, pigment production was qualitatively lower when cultured with the yeast (Fig.2B).

### Phenotyping of intestinal fungal isolates

We conducted assays to test the fungi’s resistance to temperature, oxidative stress, and bile salts to determine whether some culturable fungi were true commensals or merely passengers introduced through the diet and dispersed via feces. Commensal candidates were expected to exhibit enhanced tolerance to conditions reflective of the intestinal environment.

As expected, a progressive reduction in growth ability was observed in correspondence with incubation temperature, which was particularly relevant for filamentous fungi (Fig.3). All yeasts could thrive at temperatures ranging from 29°C to 37°C. In contrast, the growth of *R. mucilaginosa* strains was impeded at 42°C (Fig.3A). At the same time, most of the yeasts tolerated bile salts; *R. mucilaginosa* demonstrated only limited proliferation in media supplemented with 2 % taurocholate (Fig. 3A). In stress-tolerance assays, *L. elongisporus* proved more susceptible than *C. albicans* and *C. tropicalis* to bile acids, hydrogen peroxide, and pH fluctuations. These findings suggest differential adaptations among gut mycobiota, with particular species potentially better endowed for persistence in the intestinal lumen.

**Figure 3.**
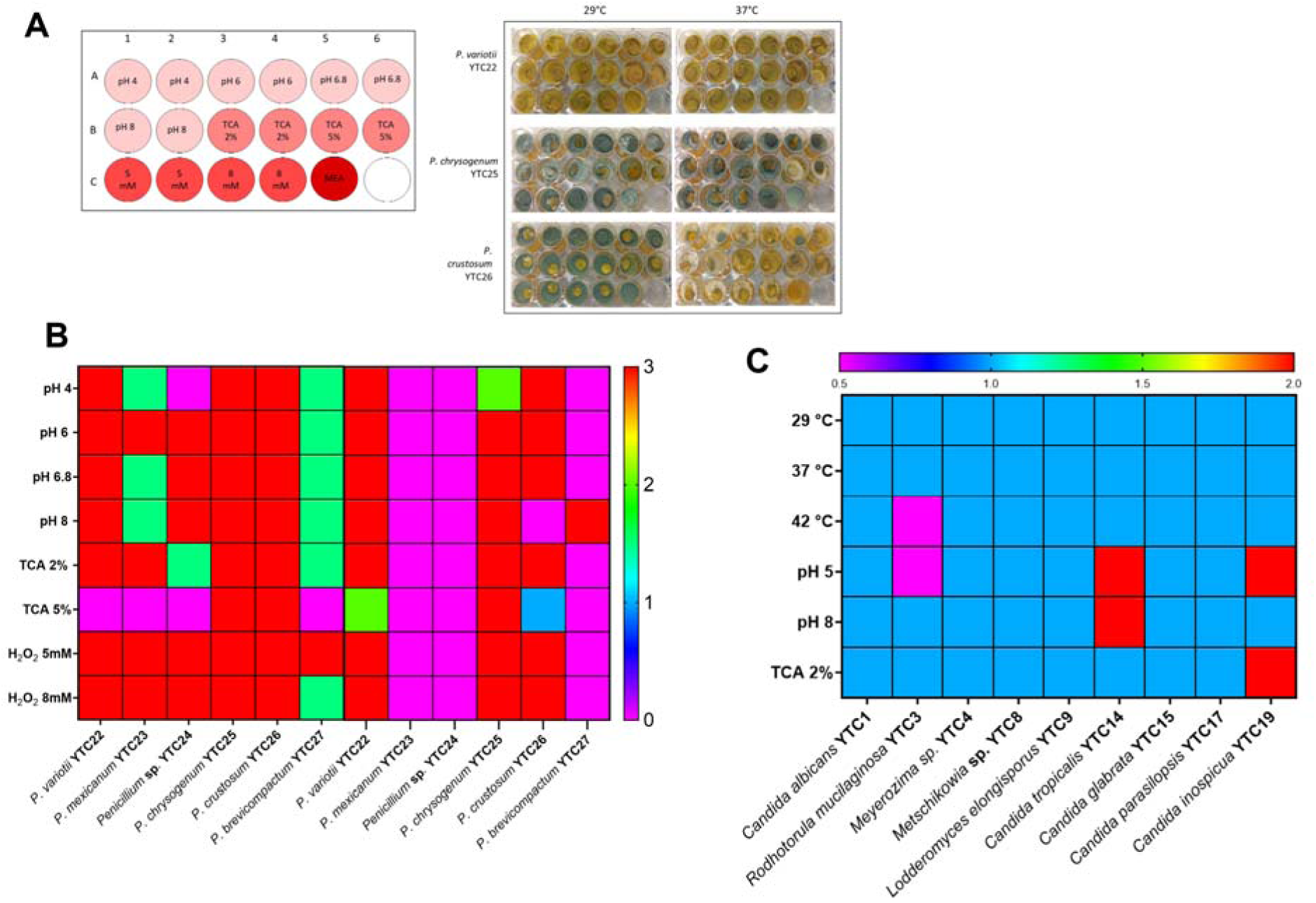
*In vitro* fungal growth and thermotolerance. **A.** Experimental plate design and illustrative growth of filamentous fungi under tested conditions. **B.** Filamentous fungi growth under different conditions. *P. variotii* YTC22 was not affected by the conditions tested. YTC25 pigment production was affected by 5% taurocholate, while 37° C severely impaired YTC26 pigmentation growth. **C** Heatmap of yeast growing under different conditions. All yeasts are thermotolerant, although R. mucilaginosa showed reduced growth at 42°C and pH 5. TCA taurocholic acid

While most environmental fungi cannot withstand mammalian body temperatures, recent years have seen the emergence of novel fungal pathogens, driven in part by the evolution of thermotolerance. Given that *P. variotii* is both thermotolerant and an opportunistic human pathogen, we assessed its competitive interactions with other filamentous isolates using a dual-culture confrontation assay on PDA. In pairwise cocultures with *P. variotii* YTC22, this strain generally outgrew its competitors, except for *P. crustosum* YTC26, which markedly inhibited its expansion (Fig. 4A).

**Figure 4.**
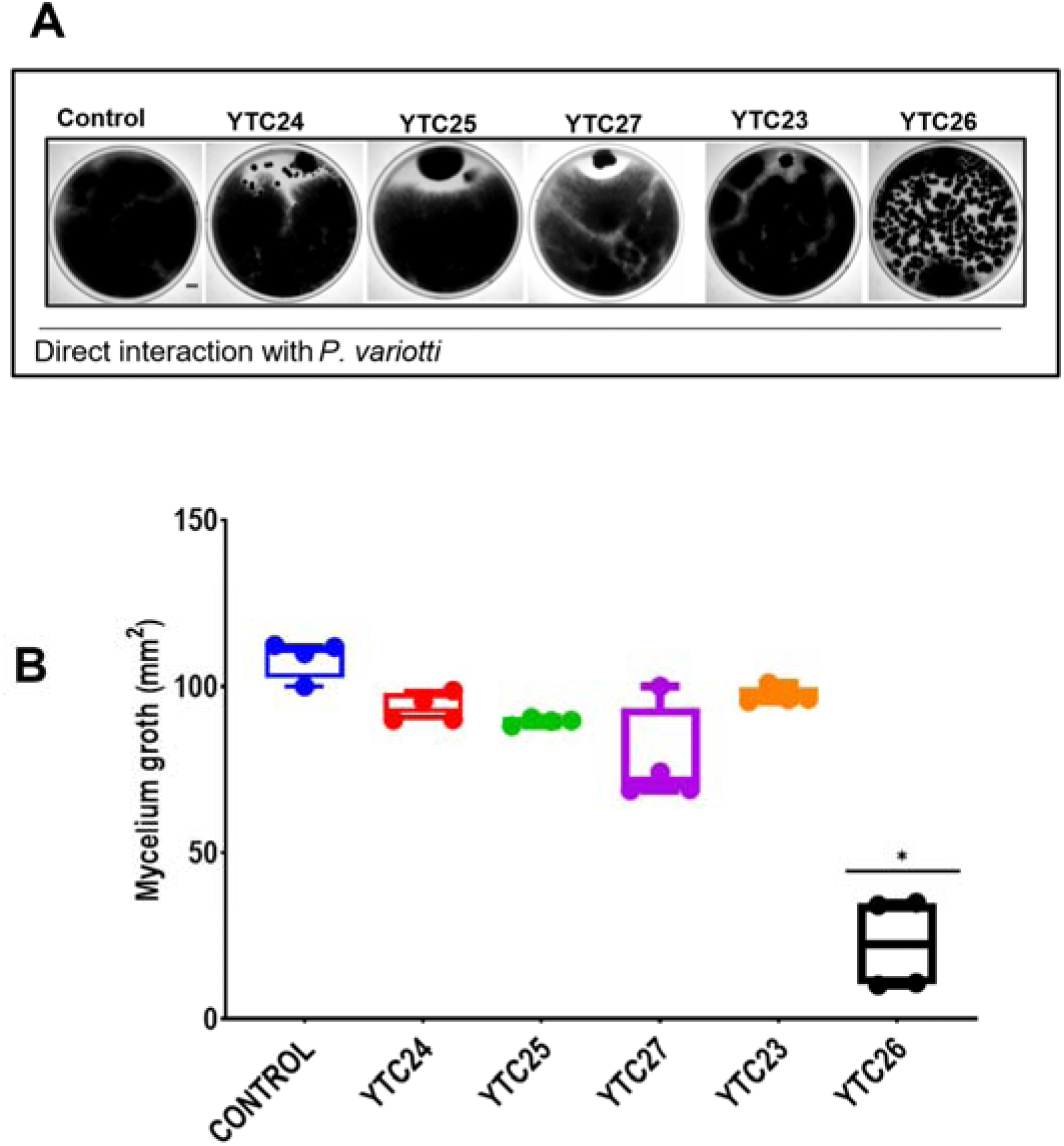
Confrontation assays against *P. variotii*. Fungi were inoculated into PDA medium at one extreme of the Petri dish and incubated at 24°C for 72 h. *P. variotti* inhibited the tested fungi, but YTC26 inhibited the *P. variotii* growth in confrontation (A). Representative images of the interaction assay are included for visual illustration. (B) Growth was evaluated by measuring the mycelial diameter on the dish relative to the control (*P. variotii* growth alone). Data represent the mean [ ± SD] of three independent experiments. Asterix indicates a statistically significant difference, according to a one-way analysis of variance [ANOVA] [p ≤ 0.05] followed by the Tukey test.

### Cytotoxic activity against Caco-2 Cells

To assess the cytotoxic potential of fungal metabolites from human feces, we performed MTT assays on Caco-2 cells using cell-free supernatants (CFS; 5, 10, 15, and 20 µL) and organic extracts (20 and 200 µg/mL). CFS were collected from ten yeast strains (YTC1, YTC3, YTC4, YTC8, YTC9, YTC10, YTC14, YTC15, YTC17, and YTC19) and three filamentous isolates (YTC22, YTC25, and YTC26). After 24 h of exposure to yeast CFS, only 20 µL of *C. inopsicua* CFS marginally reduced cell viability (Fig. 5A). In contrast, CFS from the filamentous fungi decreased viability by approximately 30% relative to controls (Fig. 5B). Consequently, we evaluated the cytotoxicity of organic extracts from YTC22, YTC25, and YTC26 at 20 and 200 µg/mL, observing moderate toxicity for the *P. variotii* extract at 200 µg/mL (Fig. 5C). To further define its concentration-response profile, *P. variotii* extracts were tested over a concentration series ranging from 0.4 µg/mL to 200 µg/mL (Fig. 5D).

**Figure 5.**
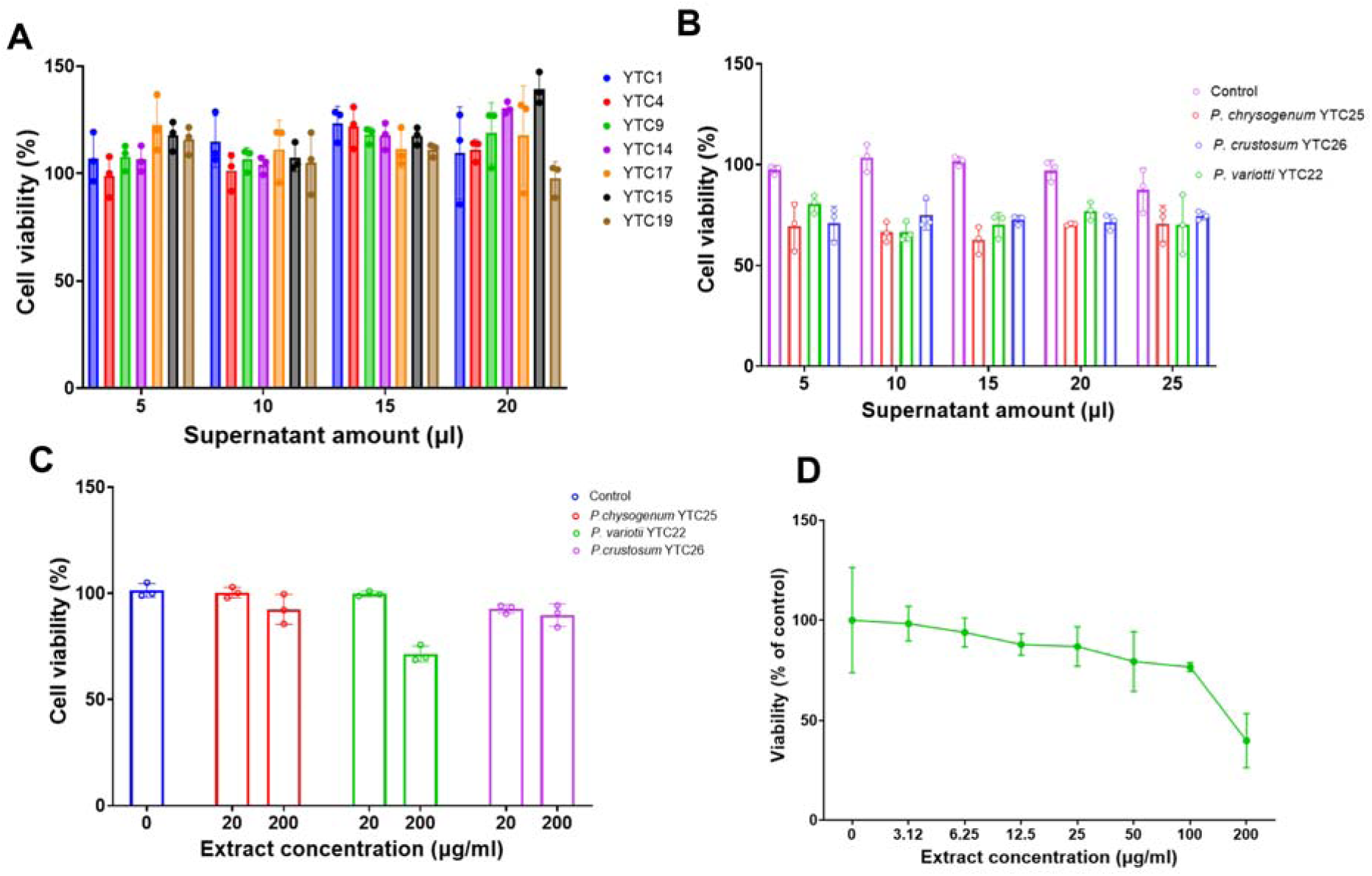
Cytotoxicity of CFS and organic extracts of thermotolerant fungi against the Caco-2 cancer cell line. (A). The cytotoxic activities of cell-free supernatants of selected fecal yeasts against Caco-2 cells. Yeasts were grown in PDA and incubated at 37°C for 24 h. (B). The cytotoxic activities of cell-free supernatants of thermotolerant molds against Caco-2 cells. Fungi were cultivated in MEA at 37°C for 7 days. (C). Cytotoxicity of organic extracts from thermotolerant fungi was tested using the MTT assay at 20 and 200 µg/mL. (D) Concentration-response curve of Caco-2 cells after being treated with organic extracts of P. variotii YTC22.

### Metabolomics analysis

Emerging evidence indicates that the gut mycobiota contributes to intestinal homeostasis and pathology by producing diverse metabolites. Both endogenous fungal metabolites and those derived from dietary fungi may act as bioactive signals, yet their roles remain poorly characterized. Therefore, we performed a dereplication analysis and untargeted metabolomic profiling on three thermotolerant filamentous fungi isolated from human feces to elucidate their metabolic repertoires.

The metabolic profile of each fungus during aerobic growth showed distinct patterns of molecule production (Table 1). *Paecilomyces variotii* YTC22 metabolite features observed in the HRMS-MS/MS data were arranged in a molecular network of 369 nodes, grouped into 20 clusters (3 nodes per cluster, 23 with 2 nodes, and 164 singletons). Dereplication analysis using the *in house* database of fungal mycotoxins revealed the presence of the mycotoxins *S*-sydonol, altenuene fellutamide B, neosartorin, ascosalipyrone, 6-[(3*E*,5*E*,7*S*)-5,7-dimethyl-2-oxonona-3,5-dienyl]-2,4-dihydroxy-3-methylbenzaldehyde, misakimycin, fonsecin, chochilioquinone D, malettinin B, and chlamydospordiol, while by GNPS analysis only oxaline was detected (Fig. 6). On the other hand, the molecular networking of *P. chrysogenum* YTC25 showed the metabolite features arranged in 167 nodes grouped into seven clusters with ≥3 nodes per cluster, six with two nodes, and 95 singletons (Supporting information). The metabolites talarolutin B and phomopsidin MK8383 were the only compounds detected by the dereplication analysis. Finally, the molecular network of P. crustosum YTC26 consisted of 344 nodes, grouped into 21 clusters: 3 nodes per cluster, 18 with 2 nodes, and 132 singletons (Supporting information). The *in-house* analysis identified the mycotoxins meleagrin, trichothecin, paxilline, hypothemycin, and roquefortine C, as annotated by GNPS. The annotated secondary metabolites implicated in dysbiosis-related pathologies are shown in Table 2.

**Figure 6.**
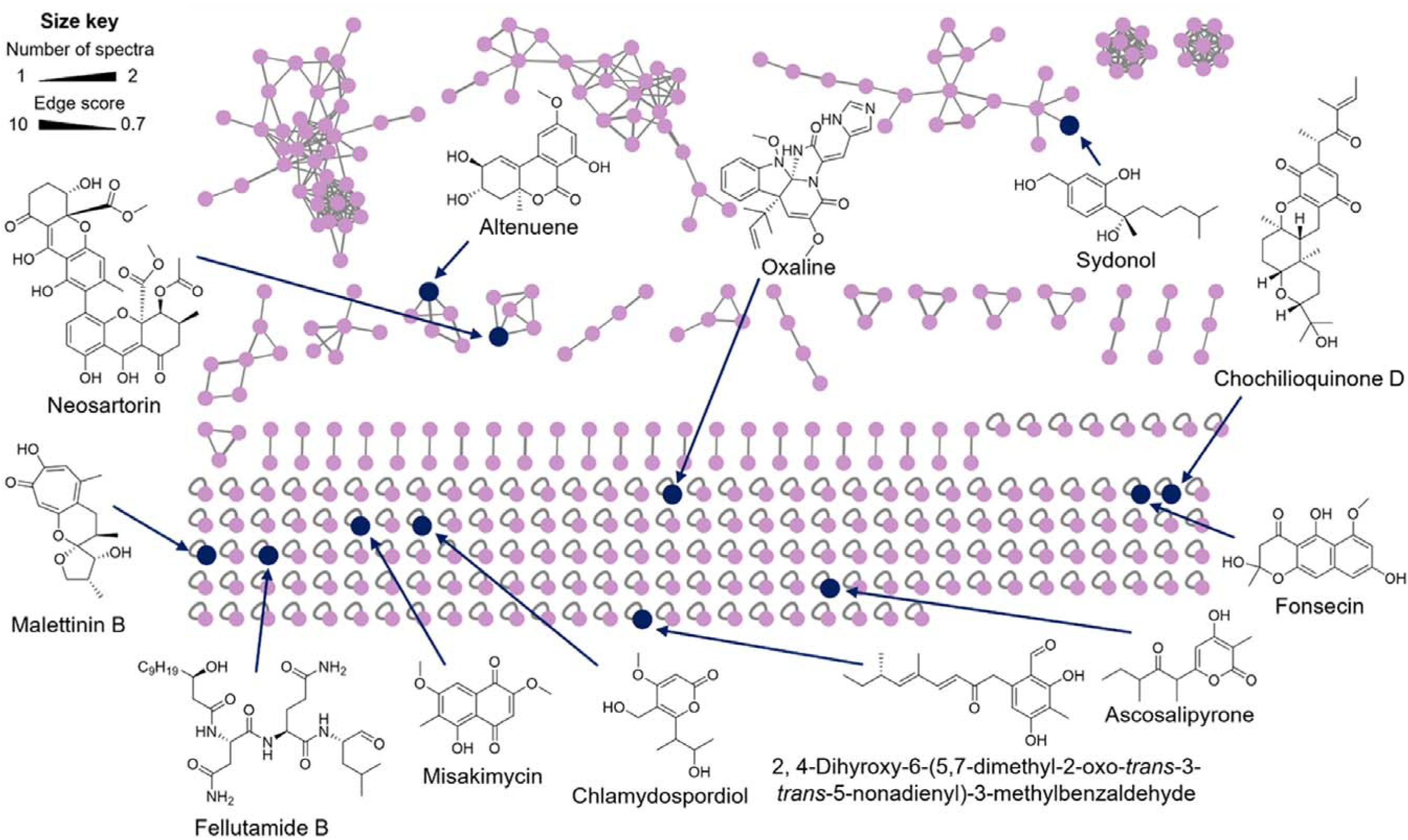
Molecular networking of *the P. variotii YTC2* organic extract, showing compounds annotated by dereplication and GNPS analysis.

**Table 1.** Chemical annotation of metabolites in *P. variotii* YTC2, *P. chrysogenum* YTC25, and *P. crustosum* YTC26 organic extract by dereplication using the *in-house* database of fungal mycotoxins and GNPS analysis.

| No. | Compound | Observed ion (*m/z*)<sup>a</sup> | Adduct | Molecular formula of the adduct | Exact Mass (calculated) | Mass accuracy (ppm) |
| --- | --- | --- | --- | --- | --- | --- |
| ***Paecilomyces variotii* YTC22** |  |  |  |  |  |  |
| 1 | S-Sydonol<sup>a</sup> | 235.1697 | [M+H–H<sub>2</sub>O]<sup>+</sup> | C<sub>15</sub>H<sub>23</sub>O<sub>2</sub> | 235.1693 | +1.9 |
| 2 | Altenuene<sup>a</sup> | 293.1027 | [M+H]<sup>+</sup> | C<sub>15</sub>H<sub>17</sub>O<sub>6</sub> | 293.1020 | +2.5 |
| 3 | Fellutamide B<sup>a</sup> | 556.3710 | [M+H]<sup>+</sup> | C<sub>27</sub>H<sub>50</sub>N<sub>5</sub>O<sub>7</sub> | 556.3705 | +0.9 |
| 4 | Neosartorin<sup>a</sup> | 681.1824 | [M+H]<sup>+</sup> | C<sub>34</sub>H<sub>33</sub>O<sub>15</sub> | 681.1814 | +1.5 |
| 5 | Ascosalipyrone<sup>a</sup> | 239.1284 | [M+H]<sup>+</sup> | C<sub>13</sub>H<sub>19</sub>O<sub>4</sub> | 239.1278 | +2.6 |
| 6 | 6-[(3*E*,5*E*,7*S*)-5,7-Dimethyl-2-oxonona-3,5-dienyl]-2,4-dihydroxy-3-methylbenzaldehyde<sup>a</sup> | 317.1755 | [M+H]<sup>+</sup> | C<sub>19</sub>H<sub>25</sub>O<sub>4</sub> | 317.1747 | +2.4 |
| 7 | Misakimycin<sup>a</sup> | 277.0712 | [M+H+CO]<sup>+</sup> | C<sub>14</sub>H<sub>13</sub>O<sub>6</sub> | 277.0707 | +2.4 |
| 8 | Fonsecin<sup>a</sup> | 291.0871 | [M+H]<sup>+</sup> | C<sub>15</sub>H<sub>15</sub>O<sub>6</sub> | 291.0863 | +2.7 |
| 9 | Chochilioquinone D<sup>a</sup> | 471.2746 | [M+H]<sup>+</sup> | C<sub>28</sub>H<sub>39</sub>O<sub>6</sub> | 471.2741 | +1.0 |
| 10 | Malettinin B<sup>a</sup> | 293.1392 | [M+H]<sup>+</sup> | C<sub>16</sub>H<sub>21</sub>O<sub>5</sub> | 293.1384 | +2.4 |
| 11 | Chlamydospordiol<sup>a</sup> | 229.1075 | [M+H]<sup>+</sup> | C<sub>11</sub>H<sub>16</sub>O<sub>5</sub> | 229.1071 | +2.0 |
| 12 | Oxaline<sup>b</sup> | 448.2017 | [M+H]<sup>+</sup> | C<sub>24</sub>H<sub>26</sub>N<sub>5</sub>O<sub>4</sub> | 448.1978 | +8.4 |
| ***Penicillium chrysogenum* YTC25** |  |  |  |  |  |  |
| 13 | Talarolutin B<sup>a</sup> | 365.2330 | [M+H]<sup>+</sup> | C<sub>21</sub>H<sub>33</sub>O<sub>5</sub> | 365.2323 | +2.1 |
| 14 | Phomopsidin MK8383<sup>a</sup> | 331.2275 | [M+H]<sup>+</sup> | C<sub>21</sub>H<sub>31</sub>O<sub>3</sub> | 331.2268 | +2.2 |
| ***Penicillium crustosum* YTC26** |  |  |  |  |  |  |
| 15 | Meleagrin<sup>a</sup> | 434.1815 | [M+H]<sup>+</sup> | C<sub>23</sub>H<sub>24</sub>N<sub>5</sub>O<sub>4</sub> | 434.1823 | –1.8 |
| 16 | Trichothecin<sup>a</sup> | 333.1696 | [M+H]<sup>+</sup> | C<sub>19</sub>H<sub>25</sub>O<sub>5</sub> | 333.1697 | –0.2 |
| 17 | Paxilline<sup>a</sup> | 436.2486 | [M+H]<sup>+</sup> | C<sub>27</sub>H<sub>34</sub>NO<sub>4</sub> | 436.2482 | +0.8 |
| 18 | Hypothemycin<sup>a</sup> | 379.1396 | [M+H]<sup>+</sup> | C<sub>19</sub>H<sub>23</sub>O<sub>8</sub> | 379.1387 | +2.3 |
| 19 | Roquefortine C<sup>b</sup> | 390.1953 | [M+H]<sup>+</sup> | C<sub>22</sub>H<sub>24</sub>N<sub>5</sub>O<sub>2</sub> | 390.1925 | +7.3 |

<sup>a</sup> *In-house* dereplication of fungal mycotoxins; <sup>b</sup> GNPS annotation.

**Table 2.**
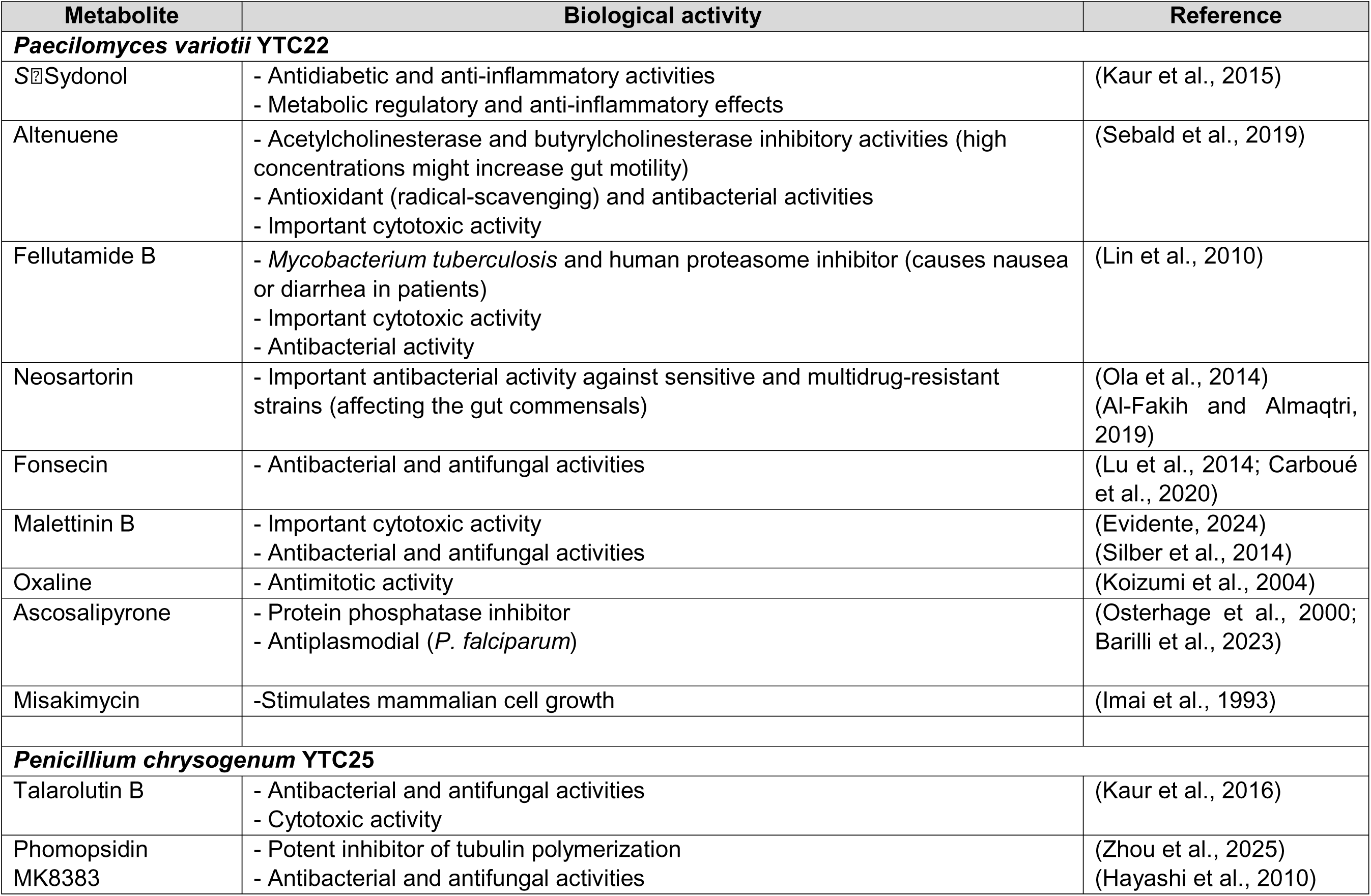
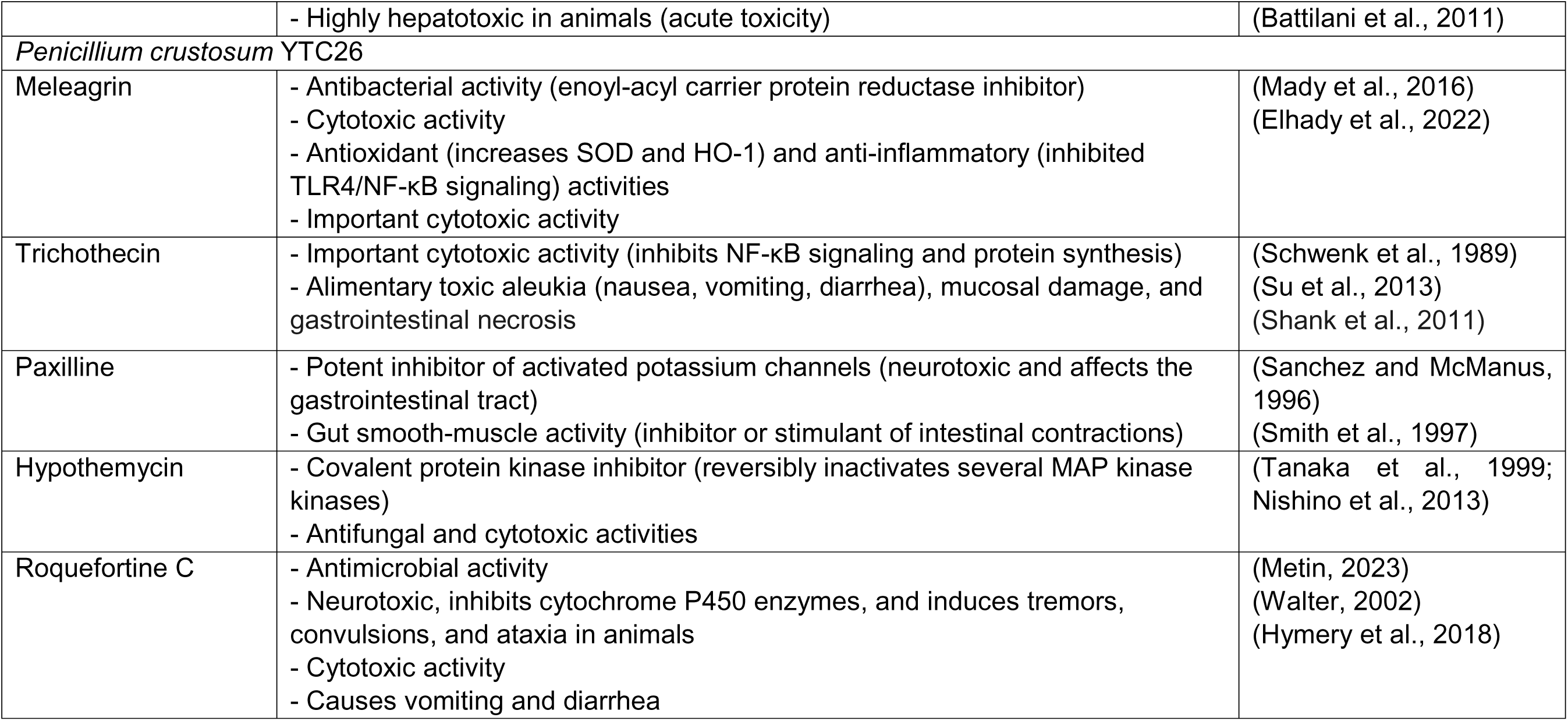
Biological activity of metabolites detected in *P. variotii* YTC22, *P. chrysogenum* YTC25, and *P. crustosum* YTC26 by dereplication and untargeted metabolomic analysis.

## Discussion

In healthy adults, the gut mycobiome is characterized by low species richness and is predominantly composed of yeast, especially members of the *Saccharomyces*, *Malassezia*, and *Candida* genera. In our cohort of patients with nosocomial diarrhea, yeasts remained dominant but with a distinctly altered composition: *Candida* and *Rhodotorula* species prevailed, and notably, no *Saccharomyces* spp. were detected. Although *C. albicans* is the archetypal core gut yeast in healthy individuals (Strati et al., 2016), environmental yeasts such as *Rhodotorula* spp., commonly transmitted via air and food, emerged as the second-most-abundant genus among our isolates (Hof, 2019). Prior work has reported intestinal *Rhodotorula* colonization in up to 5% of healthy children and 12% of young adults (Strati et al., 2016). Our data affirm its underrecognized presence in the gut ecosystem, or at least during dysbiosis. Less prevalent opportunistic yeasts, including *N. glabratus*, *C. inopsicua*, *C. parapsilosis*, and *C. tropicalis*, each accounted for approximately 5% of samples, aligning with their sporadic but clinically relevant presence in the gastrointestinal tract (Strati et al., 2016; Vargas-Espindola et al., 2023). In particular, immunocompromised patients often experience fungemia caused by these same *Candida* species (Vargas-Espindola et al., 2023).

We also detected *L. elongisporus*, a rarely reported opportunist often misidentified as *C. parapsilosis* due to phenotypic similarity (Wang and Xu, 2023). Its presence in fecal samples likely reflects antibiotic-induced dysbiosis and nosocomial exposures. Notably, we recovered putative probiotic yeasts, *Meyerozyma* and *Metschnikowia* spp., from two patients, suggesting a potentially underexplored niche for beneficial fungi in the gut ecosystem.

Filamentous fungi are relatively uncommon in both healthy and diseased adult guts. In healthy individuals, environmental and food-associated genera, such as *Cladosporium*, *Penicillium*, and *Aspergillus,* are occasionally detected, though their roles in the gut remain speculative. In contrast, our cohort yielded only *Penicillium* species as fungal isolates. Growth-temperature assays showed that two of five *Penicillium* isolates thrived at 37°C, while *P. variotii* isolates grew at 42°C, highlighting their thermotolerance. Although these molds are often dismissed as contaminants, they are increasingly implicated in invasive infections in both immunocompromised and immunocompetent patients. For instance, *P. variotii* has been associated with 59 reported human infections through 2020, exhibiting a 28.8 % overall mortality and accounting for ten direct deaths (Sprute et al., 2021).

Co-occurrence analyses revealed that *Salmonella-*positive samples harbored *L. elongisporus* and *R. mucilaginosa*, and *C. albicans* was present in samples positive with *Clostridioides difficile*, corroborating previous reports of bacterial–fungal interactions in dysbiosis (van Leeuwen et al., 2016; Lambooij et al., 2017). Although formal interkingdom association tests were beyond the scope of this study, the co-isolation of yeasts and molds from individual patients suggests potential ecological interactions within the gut niche.

Functional assays using yeast cell-free supernatants (CFS) revealed modest cytotoxicity against Caco-2 cells. Notably, CFS from *C. inopsicua* YTC19 elicited the most significant reduction in viability, while supernatants from the thermotolerant filamentous strains *P. variotii* YTC22, *P. chrysogenum* YTC25 and *P. crustosum* YTC26 decreased cell viability by approximately 40%. When tested as organic extracts, only the fraction derived from *P. variotii* YTC22 retained cytotoxic activity, suggesting that critical bioactive compounds were lost during extract processing.

Untargeted metabolomic profiling and dereplication of organic extracts from the three thermotolerant fungi revealed diverse metabolites with documented bioactivities, some of which could contribute to diarrheal diseases and dysbiosis through multifaceted mechanisms. These fungi produce metabolites with antimicrobial, cytotoxic, neuromodulatory, and immunomodulatory properties, creating a “perfect storm” of gut dysfunction when acting in concert. For example, compounds such as neosartorin, fonsecin, meleagrin, malettinin B, talarolutin B, phomopsidin MK8383, and roquefortine C display broad-spectrum antibacterial (and in some cases antifungal) effects. By targeting both Gram-positive and Gram-negative commensals, these metabolites could significantly reduce microbial diversity and decrease the production of short-chain fatty acids, which are crucial for maintaining colonic epithelial health. Concomitant suppression of colonization resistance paves the way for opportunistic pathogens such as *Clostridioides difficile* to overgrow, creating an osmotic and inflammatory environment that favors diarrheal output.

Beyond disrupting microbial communities, several of these molecules exert direct cytotoxic effects on the gut epithelium. Fellutamide B and oxaline induce apoptosis or cell-cycle arrest in rapidly dividing cells, including epithelial stem and progenitor populations. Malettinin B and hypothemycin similarly target the proliferative machinery through kinase inactivation, impairing epithelial renewal. Of particular note, the trichothecene trichothecin causes mucosal necrosis and the classic syndrome of “alimentary toxic aleukia,” in which severe diarrhea arises from direct epithelial injury and compromised barrier integrity (Schwenk et al., 1989; Su et al., 2013). Loss of barrier function increases intestinal permeability (“leaky gut”), permitting luminal antigens and microbial products such as lipopolysaccharide to drive further inflammation, fluid secretion, and diarrheal symptoms.

Gut motility, governed by enteric neurons and smooth muscle ion channels, represents a third axis of disruption by fungal metabolites. Altenuene’s inhibition of acetyl- and butyryl-cholinesterases elevates acetylcholine at neuromuscular junctions, resulting in accelerated peristalsis and osmotic diarrhea at high exposures (Sebald et al., 2019). Paxilline’s blockade of BK (maxi-K) potassium channels similarly perturbs smooth-muscle contractions, producing alternating inhibition and stimulation of intestinal transit that may clinically manifest as bouts of constipation interspersed with diarrhea (Sanchez and McManus, 1996; Smith et al., 1997). Moreover, gastrointestinal side effects such as nausea and diarrhea are well-documented in patients receiving proteasome inhibitors like fellutamide B analogs, underscoring neuronal and mucosal contributions to dysmotility (Lin et al., 2010).

Finally, immune modulation by specific metabolites can indirectly facilitate gut dysfunction. Anti-inflammatory agents such as *S*-sydonol and meleagrin inhibit NF-κB signaling and reduce cytokine production, which, when combined with antimicrobial pressure, may paradoxically impair mucosal defenses against invading pathogens(Kaur et al., 2016; Elhady et al., 2022). Ascosalipyrone’s inhibition of protein phosphatases could disrupt tight-junction assembly and skew epithelial signaling cascades, further compromising barrier integrity. Altered immune tone may diminish pathogen clearance or skew T-helper cytokine balances (Th1/Th17), contributing to subclinical inflammation, increased permeability, and secretory diarrhea.

Several key knowledge gaps remain: it is unclear whether the *in situ* concentrations of these metabolites within the gastrointestinal tract reach bioactive levels following ingestion or colonization; the interactions between the mycobiome and bacterial microbiota may modulate fungal metabolism, enhancing or attenuating the production of these compounds (i.e., have not been quantified). Alternatively, if unidentified fungal metabolites could act as inducers or repressors of bacterial toxin production, this would further complicate host–microbe interactions. Future investigations will be crucial for understanding the contributions of fungi to gastrointestinal diseases and for developing interventions that preserve gut health in the face of fungal metabolite exposure.

## Conclusions

Our analysis of hospitalized Mexican adults revealed a gut mycobiome markedly shifted toward opportunistic and thermotolerant fungi. *Candida* and *Rhodotorula* species predominated, alongside filamentous genera such as *Penicillium* and *Paecilomyces,* likely reflecting antibioticl1linduced dysbiosis and environmental exposures. The col1loccurrence of these fungi with bacterial pathogens *(Salmonella* spp. and *Clostridioides difficile*) suggests potential interkingdom interactions that merit targeted investigation. Functional assays demonstrated that yeast celll1lfree supernatants exert modest cytotoxicity on intestinal epithelial cells, with the organic extract of *P. variotii* retaining significant activity, highlighting the importance of fungal metabolites in modulating host tissues. Untargeted metabolomic profiling further identified diverse bioactive compounds that may contribute to dysbiosisl1lassociated pathologies. These findings lay the groundwork for future studies aimed at elucidating fungal-bacterial-host interactions and characterizing mycobiomel1lderived metabolites, with the ultimate goal of developing strategies to mitigate fungal pathogenicity and harness beneficial fungal functions in clinical settings.

## Acknowledgments

Romina Rodríguez-Sanoja, Silvia Moreno-Mendieta, and Enrique Ortega for generously providing reagents and laboratory space for our pilot experiments, and Carlos A. Fajardo-Hernández and Mariano Jacome for their support on the initial metabolomic analysis.

## Funding and additional information

This work was partially supported by grants from UNAM-DGAPA PAPIIT IA206823 (A.R.R.) and IN203923 (M. F.), and FQ-PAIP 5000-9145 (M. F.).

## Conflict of interest

The authors declare no conflict of interest.

## Data Availability

Sequencing data generated in this study have been deposited in the European Nucleotide Archive under BioProject PRJEB107195.

## Notes

### Competing Interest Statement

The authors have declared no competing interest.

