## SUPPLEMENTAL MATERIAL for "Isolation and metabolomic analysis of the culturable human gut mycobiota during dysbiosis"

**Supporting information**

**Figure S 1**. Maximum likelihood (ML) tree of *Paecilomyces* genus based on ITS (A) and β-tubulin sequences (B). The ITS tree is rooted with Mucor mucedo (CBS 640.67). Constructions were carried out in MEGA-X using the maximum likelihood method with 1000 bootstrap replicates and the Tamura-Nei model. Fecal fungi isolated in this present work are shown in red.

**Figure S 2** Maximum likelihood (ML) tree of *Penicillium* genus based on ITS (A) and β-tubulin sequences (B). The ITS tree is rooted with *Mucor mucedo* (CBS 640.67). Constructions were carried out in MEGA-X with the maximum likelihood method with bootstrap 1000 and the Tamura-Nei model. Fecal fungi isolated in this present work are shown in red*.*

**Figure S 3**. Maximum likelihood (ML) tree of *Candida* (A) and *Rhodotorula* (B) genera based on ITS sequences. The Candida ITS tree includes R. mucilaginosa, and the Rhodotorula genus tree is rooted with *Candida albicans* (CBS 640.67). Constructions were carried out in MEGA-X using the maximum likelihood method with 1000 bootstrap replicates and the Tamura-Nei model. Fecal fungi isolated in this present work are shown in red.

**Figure S 4**. Maximum likelihood (ML) tree of yeast based on ITS sequences. The tree is rooted with *R. mucilaginosa* (CBS 316). Constructions were carried out in MEGA-X using the maximum likelihood method with bootstrap 1000 and the Tamura-Nei model. Fecal fungi isolated in this present work are shown in red.

**Figure S 5** Molecular networking of *P. chrysogenum* YTC25 organic extract showing the compounds annotated by dereplication and GNPS analysis.

**Figure S 6**. Molecular networking of *P. crustosum* YTC26 organic extract showing the compounds annotated by dereplication and GNPS analysis.

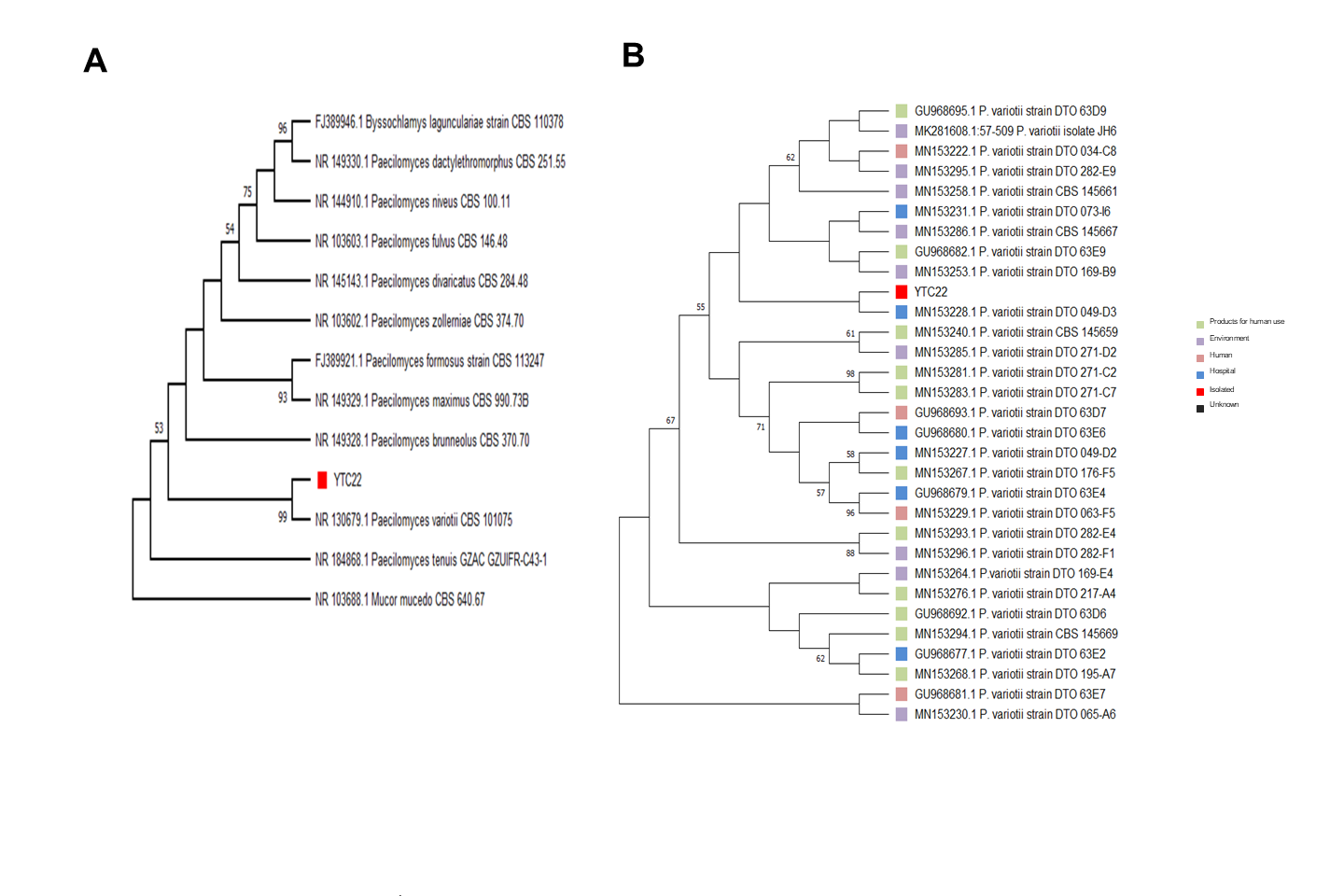

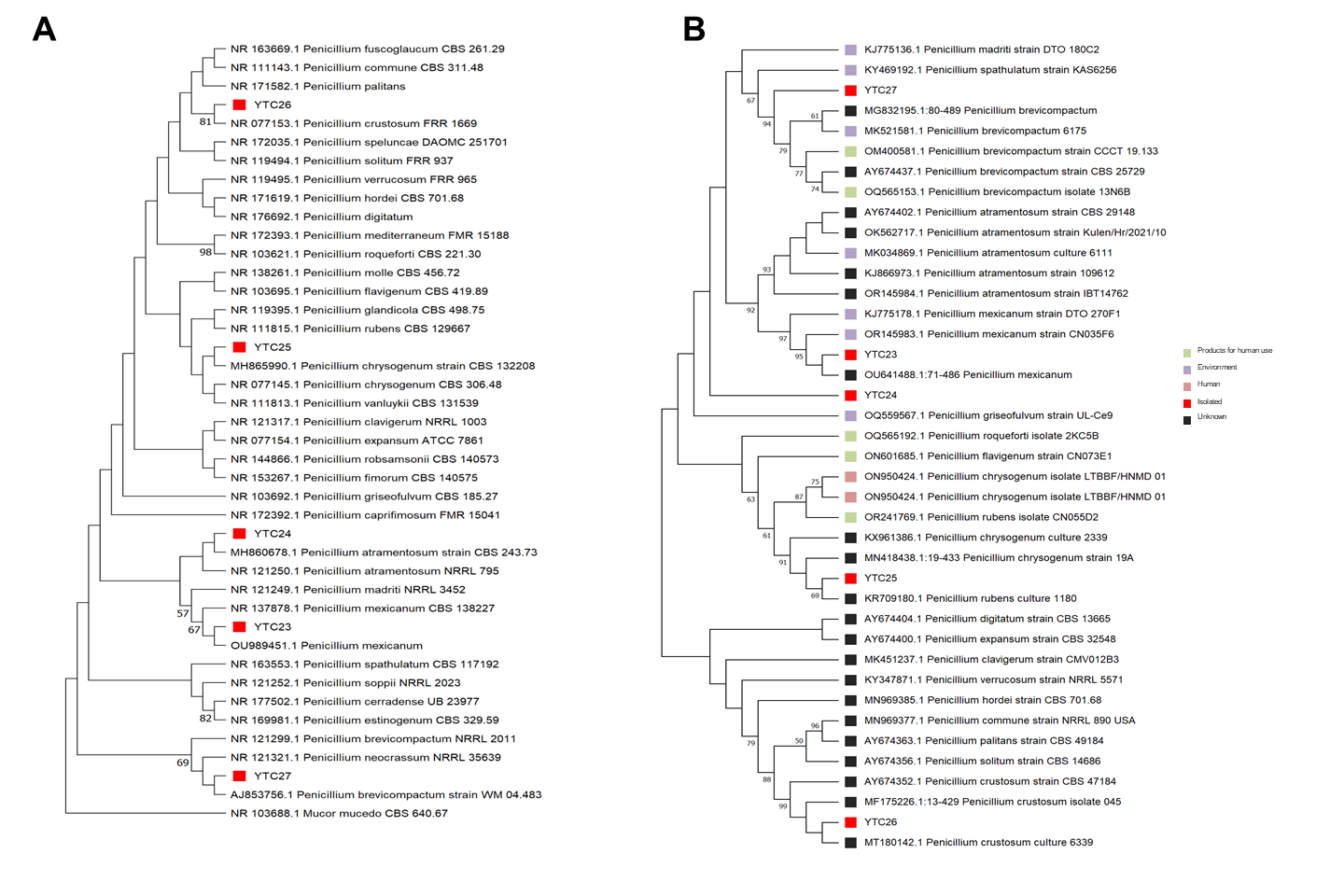

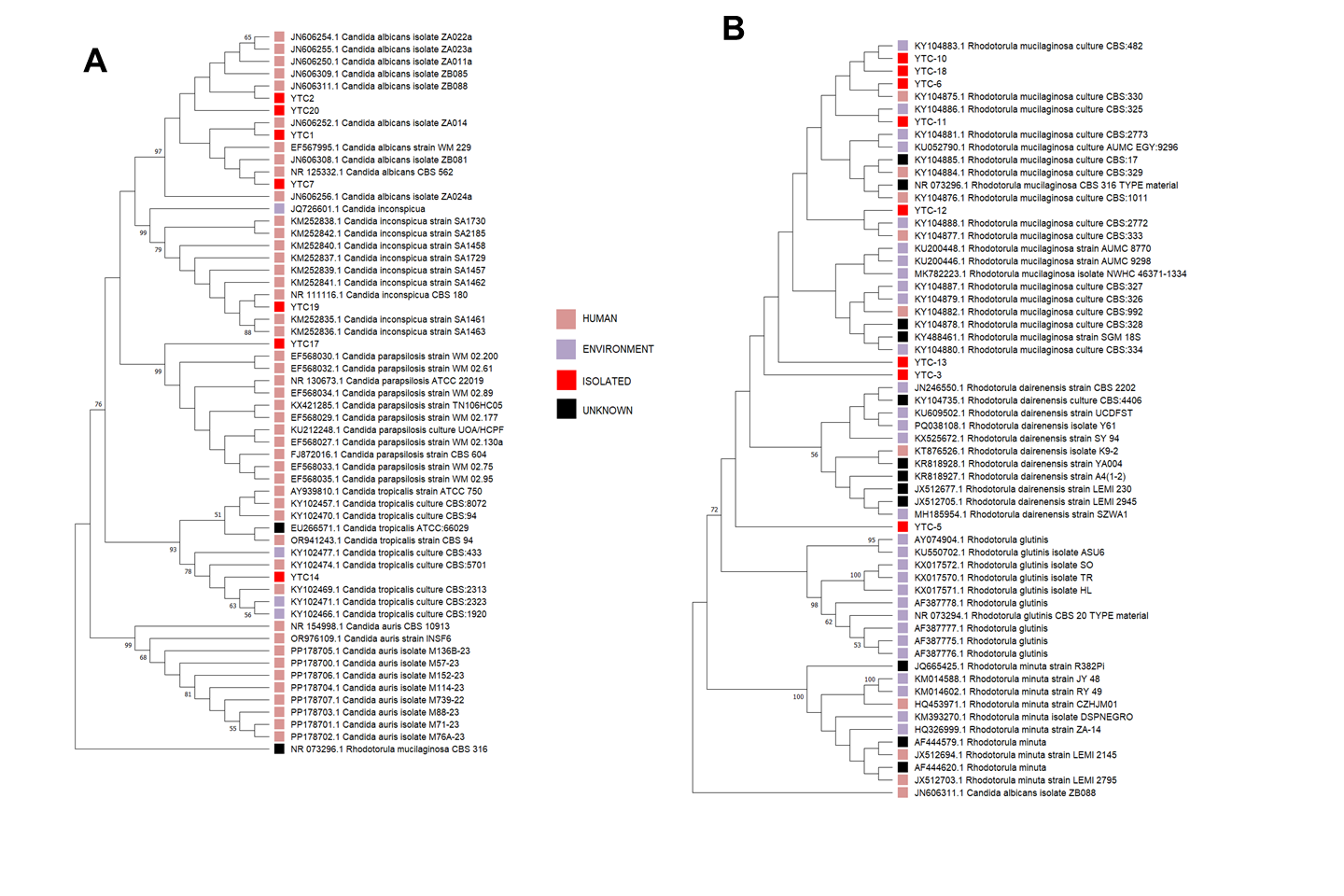

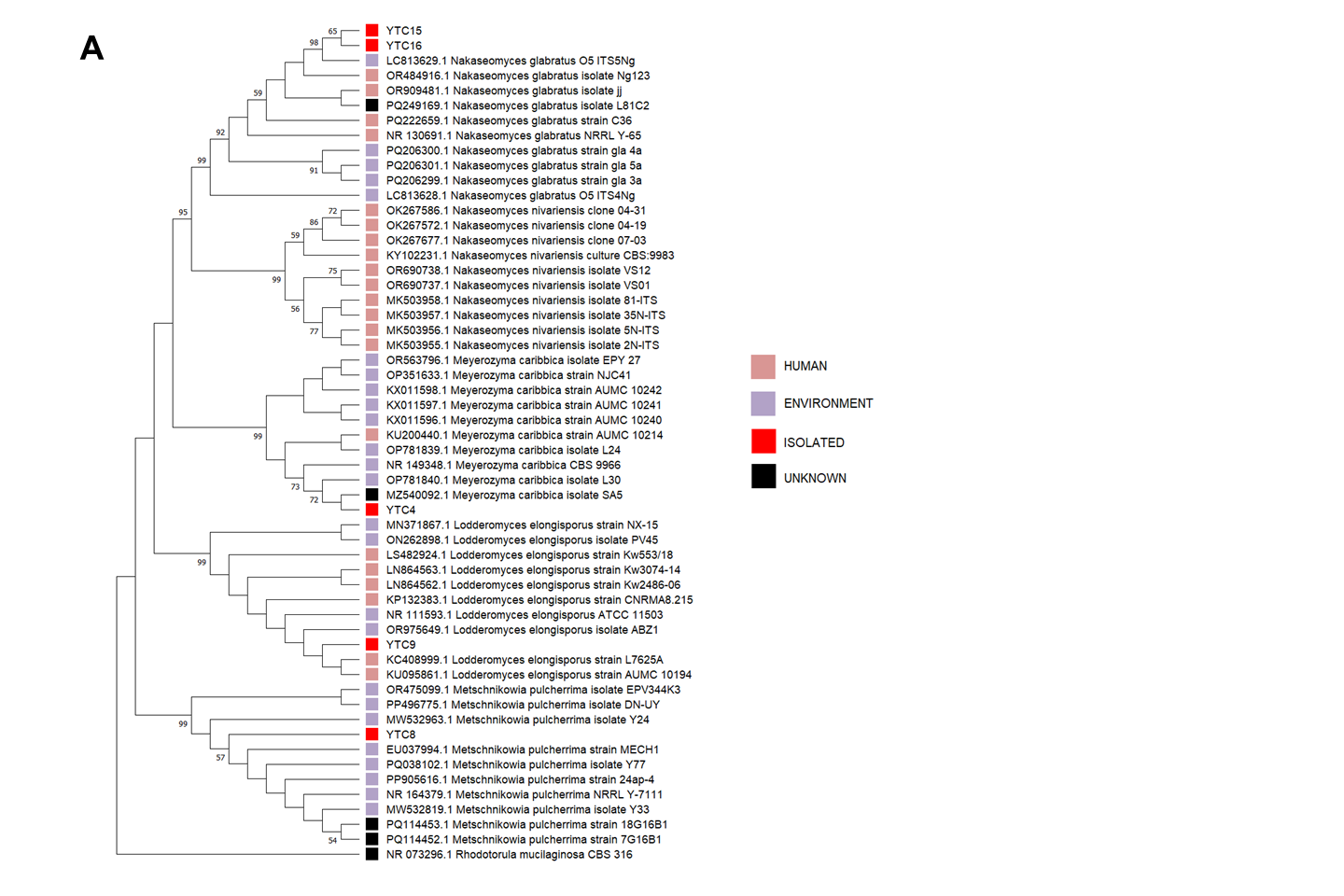

**Figure S5.** Molecular networking of *P. chrysogenum* YTC25 organic extract showing the compounds annotated by dereplication and GNPS analysis.

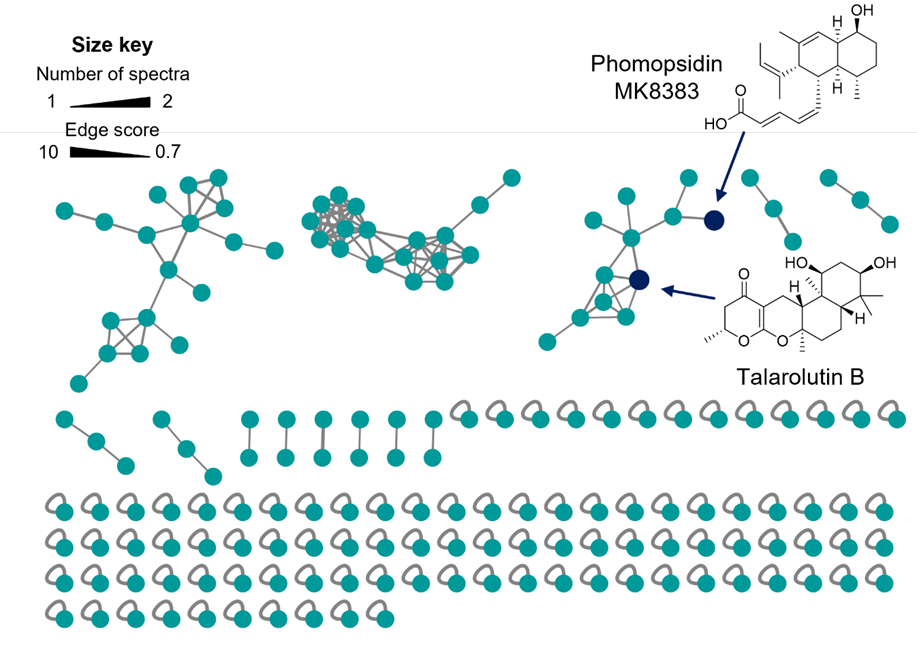

**Figure S6.** Molecular networking of *P. crustosum* YTC26 organic extract showing the compounds annotated by dereplication and GNPS analysis.

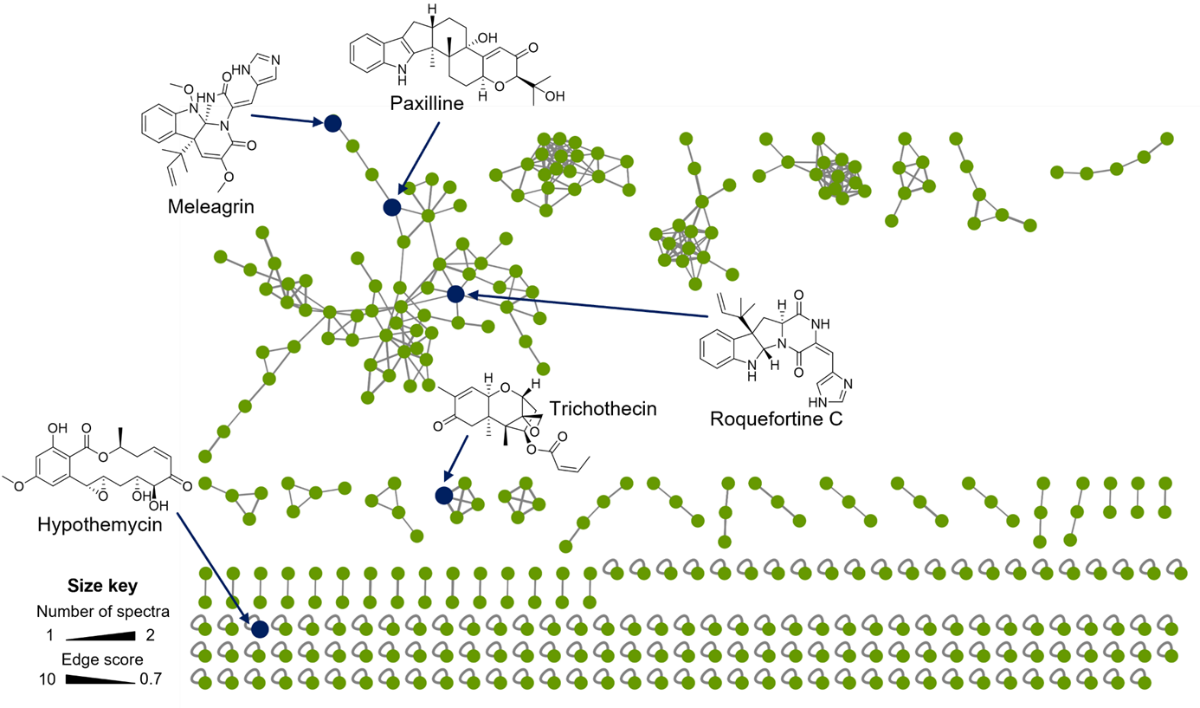
